# AITeQ: A machine learning framework for Alzheimer’s prediction using a distinctive 5-gene signature

**DOI:** 10.1101/2023.09.16.558056

**Authors:** Ishtiaque Ahammad, Anika Bushra Lamisa, Arittra Bhattacharjee, Tabassum Binte Jamal, Md. Shamsul Arefin, Zeshan Mahmud Chowdhury, Mohammad Uzzal Hossain, Keshob Chandra Das, Chaman Ara Keya, Md Salimullah

## Abstract

Neurodegenerative diseases, such as Alzheimer’s disease, pose a significant global health challenge with their complex etiology and elusive biomarkers. In this study, we developed the Alzheimer’s Identification Tool using RNA-Seq (AITeQ), a machine learning model based on an optimized random forest algorithm for identification of Alzheimer’s from RNA-Seq data. Analysis of RNA-Seq data from 433 individuals, including 293 Alzheimer’s patients and 140 controls led to the discovery of 47,929 differentially expressed genes. This was followed by a machine learning protocol involving feature selection, model training, performance evaluation, and hyperparameter tuning. The feature selection process undertaken in this study, employing a combination of 4 different methodologies, culminated in the identification of a compact yet impactful set of 5 genes. Ten diverse machine learning models were trained and tested using these 5 genes (*ITGA10, CXCR4, ADCYAP1, SLC6A12, VGF*). Performance metrics, including precision, recall, F1-score, accuracy, receiver operating characteristic area under the curve, and confusion matrices, were assessed before and after hyperparameter tuning. Overall, the random forest model with optimized hyperparameters was identified as the best and was used to develop AITeQ. AITeQ is available at: https://github.com/ishtiaque-ahammad/AITeQ

**Key Points:**

- A set of 5 genes (*ITGA10, CXCR4, ADCYAP1, SLC6A12, VGF*) were identified following differential gene expression and feature importance analysis.
- Ten diverse machine learning algorithms were trained and tested using the gene expression patterns of the identified 5 genes. The random forest algorithm with customized hyperparameters was found to be the best-performing model for differentiating Alzheimer’s disease samples from control.
- AITeQ, a user-friendly, reliable, and accurate machine learning framework for Alzheimer’s disease prediction was developed based on the 5 gene signature.

## Introduction

Millions across the world are affected by Alzheimer’s disease (AD) which leads to cognitive impairments. For timely and effective treatment, early and accurate diagnosis of the disease is very important. Traditional diagnostic approaches often rely on clinical symptoms and neuroimaging, which might not capture the molecular intricacies of the disease. RNA-Seq, a high-throughput sequencing technique, offers a comprehensive snapshot of the transcriptome, enabling the identification of gene expression alterations associated with neurodegeneration [1].

Machine learning (ML) algorithms have demonstrated remarkable potential in analyzing large-scale, complex datasets like RNA-Seq data. By integrating ML techniques, researchers can identify disease-specific gene expression signatures, classify patient samples, and predict disease progression [2]. ML models learn from patterns within the data and can uncover subtle relationships that might elude traditional statistical methods. Selection of important genes from RNA-Seq data is an application of supervised ML techniques [3].

Identifying reliable biomarkers is a critical step in disease diagnosis and prognosis. ML models can aid in the discovery of potential biomarkers by pinpointing genes consistently associated with disease states. Since numerous RNA-Seq studies are based on the comparison between cases and controls, one such study focused on the development of a logistic regression model where disease state was described as a function of RNA-Seq reads [4]. The Support Vector Machine was also used for the early detection for both prediction and classification of AD [5]. Another study revealed the efficacy of the Decision tree algorithm for construction of classifiers that can classify different AD genes [6]. Random forest model was also implemented to predict the individualized conversion from mild cognitive impairment stage to AD [7]. A more robust multi stage classifier-based approach consisting of K-nearest neighbor, Support Vector Machine, and Naive Bayes classifier was reported to be able to efficiently classify AD [8]. For biomarker based early diagnosis of AD with high classification accuracy, gradient boosting algorithm was also used [9]. Analyzing single-cell RNA-Seq data from patients with AD and healthy individuals using Extreme Gradient Boosting revealed genes with diagnostic potential [10]. A meta-analysis and ML based integrative study identified differentially expressed microRNAs in blood as potential biomarkers for AD using adaptive boosting [11]. Light Gradient Boosting Machine was used for feature selection to detect AD from circulating non-coding RNAs [12].

In light of these advancements, we aimed to analyze publicly available AD-associated gene expression data and build a gene signature-based ML framework that can differentiate AD from control. For this purpose, several sophisticated feature selection methods and ML algorithms were utilized following the identification of DEGs. Findings from this study is likely to contribute to the better understanding of the genes most crucial for AD and utilize them as biomarkers.

## Methods

A visual representation of the workflow followed in the study is illustrated in **Figure 1**.

**Figure 1:**
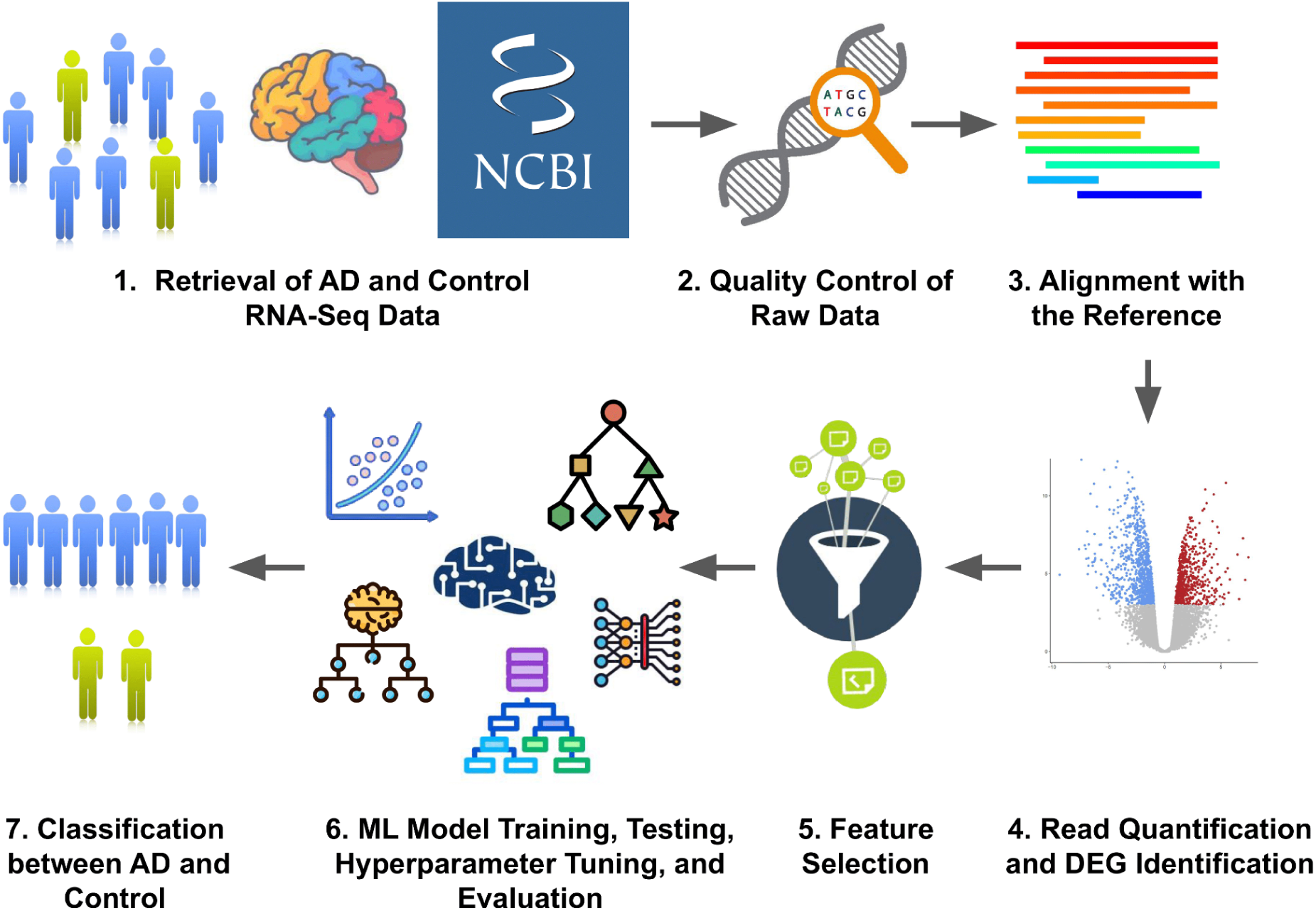
Workflow of the study. RNA-Seq data of AD and control was retrieved from NCBI. The raw reads were subjected to quality control using FastQC, and subsequently aligned with the human reference genome (GRCh38.p13) using HISAT2. The quantification of reads was performed using the featureCounts algorithm, while the identification of DEGs was conducted using the DESeq2 statistical tool. Feature selection was carried out using four methods. It was followed by 10 ML model training, testing, hyperparameter tuning, and evaluation.

### Data Retrieval and Preprocessing

The NCBI Gene Expression Omnibus (GEO) (https://www.ncbi.nlm.nih.gov/geo/) database was used to obtain the RNA-Seq datasets from 9 projects. The NCBI BioProject IDs and the sample source for these projects have been summarized in **Table 1**. Combining all datasets, the total number of samples were 433 individuals, of whom 293 were diagnosed with AD and 140 were healthy controls. The RNA-Seq data analysis workflow consisted of several steps. At first, the raw read quality was checked using FastQC (https://www.bioinformatics.babraham.ac.uk/projects/fastqc/). Next, the alignment of the reads to the *Homo sapiens* GRCh38.p13 reference genome was carried out using HISAT2 [13]. The mapped reads were then distributed to genomic features. Finally, gene expression was quantified using FeatureCounts [14]. The DESeq2 statistical tool was utilized to identify differentially expressed genes (DEGs). In order to adjust the p-values and ascertain the reliability of the identified DEGs, the False Discovery Rate (FDR) method was employed. Between the control and AD groups, the fold change (FC) of each gene was calculated. Genes with a p-adjusted value of <0.05 and a Log2FC value >1 were considered as significant DEGs [15]. The normalized and variant stabilized count of these significant DEGs were used as the features for ML.

**Table 1:** List of retrieved NCBI BioProjects and their respective sample sources.

| NCBI BioProject ID | Source |
| --- | --- |
| PRJNA675355 | Putamen |
| PRJNA683625 | Neural progenitor cells |
| PRJNA767074 | Hippocampus |
| PRJNA796229 | Substantia nigra, parietal lobe, Hippocampus, basal ganglia |
| PRJNA279526 | Hippocampus |
| PRJNA232669 | Dorsolateral prefrontal cortex |
| PRJNA377568 | Fusiform gyrus |
| PRJNA413568 | Lateral temporal lobe |
| PRJNA516886 | Fusiform gyrus |

### Feature Selection for Machine Learning Models

This study utilized a comprehensive array of feature selection strategies to unravel the most important features needed for training various ML models. The determination of feature importance was conducted through the application of four separate methodologies, namely Random Forest Classifier [16], Gradient Boosting Classifier [17], Recursive Feature Elimination (RFE) [18], and LassoCV [19]. In our study, we utilized the “feature_importances_” function of the RandomForestClassifier and GradientBoostingClassifier algorithms to assess the relative importance of each feature in the model. The Recursive Feature Elimination technique entails iteratively eliminating features with the least significance by employing a linear regression model. Furthermore, the LassoCV technique employed a Lasso linear regression model to award significance scores to features according to their coefficients. These strategies, when used together, enabled the identification of important features from our dataset. A Venn diagram was constructed with the top 10 features identified by each approach, and the set of features that were found to be common to all methods were selected. Subsequently, the selected features were utilized to build and refine ML models for Alzheimer’s disease classification.

### Machine Learning Model Training

Scaling of features is an important part of data preprocessing in most ML methodologies. In this study, the input features were scaled utilizing the “StandardScaler” function from the preprocessing module in the scikit-learn toolkit [20]. After applying feature scaling, the data was split into training dataset and testing dataset through the use of the “train_test_split” function which is available in the scikit-learn library. Eighty percent of the data constituted the training dataset, while the remaining twenty percent constituted the test dataset. The test dataset was utilized to evaluate the performance of the models that were trained on the training dataset. The training process involved the utilization of 10 ML models, namely logistic regression, support vector machine, decision tree, random forest, naive bayes, k-nearest neighbor, gradient boosting, adaptive boosting, extreme gradient boosting, and light gradient boosting machine.

### Logistic Regression

In ML, Logistic Regression is an algorithm that is frequently used for solving regression tasks where the dependent variable is categorical in nature. It predicts the probability of the dependent variables by estimating the coefficients of the independent variables in the ML model [21].

### Support Vector Machine

Support Vector Machine (SVM) is a powerful ML model which is used in both classification and regression domains. Recognition of the hyperplane that achieves the maximum separation between two classes is the primary goal of SVM. Identification of such hyperplanes relies upon the identification of the support vectors [22].

### Decision tree

Decision tree is an ML model where each internal node of the tree is equivalent to a choice made based on a particular attribute, and each leaf node corresponds to an output of classification or regression. The algorithm iteratively divides the dataset into smaller subsets. It continues to look for the feature that contains the most significant information, until a predetermined output is found [23].

### Random Forest

Random Forest is a notable ensemble learning strategy that is utilized for not just classification and regression but also feature selection. In case of ensemble learning, numerous decision trees are put to use for enhanced accuracy and generalization [16].

### Naive bayes

Naive Bayes is a Bayes’ theorem-based probabilistic model that calculates the likelihood of a class from a given set of features. It assumes that the features are independent of each other while assigning them a class label, thereby getting the name "naive" [24].

### K-nearest neighbor

K-nearest neighbors (KNN) is a non-parametric method that is mainly used to decipher problems involving classification and regression. KNN functions through the identification of neighboring data points based on their similarity [25].

### Gradient Boosting

Gradient Boosting exhibits remarkable efficacy in making predictions from intricate datasets, such as RNA-Seq data. It is an ensemble method that iteratively combines numerous weak learners in order to generate strong learners which can eventually make accurate predictions [26].

### Adaptive Boosting

The Adaptive Boosting algorithm takes an iterative approach to modify the weights assigned to the training data, with the objective of focusing on the misclassified cases in each iteration. During each iteration, Adaboost uses a weak learner to train on a certain subset of the training data. It takes into account its classification error and assigns a weight to each training example. The weights assigned to misclassified examples are augmented, while the weights assigned to correctly classified examples are diminished. This technique is iteratively implemented until a predetermined outcome is satisfied [27].

### Extreme Gradient Boosting

Extreme Gradient Boosting (XGBoost) algorithm enhances the conventional gradient boosting approach by integrating well established regularization methods such as L1 and L2 regularization, to minimize the possibility of model overfitting. Additionally, XGBoost employs a novel approach to estimate the second-order gradient of the loss function. Thus it enhances both the speed of convergence and the accuracy of the model to solve regression and classification tasks [28].

### Light Gradient Boosting Machine

The Light Gradient Boosting Machine (LightGBM) is a framework that utilizes a collection of weak learners, most commonly in the form of decision trees, with the objective of constructing a strong learner. It operates by iteratively including additional models into its ensemble learning approach, with a primary objective of lowering the gradient of the loss function [29].

### Machine Learning Model Performance Evaluation

Cross-validation is an important method in ML as it provides a more reliable estimate of the success of the model on unseen data as opposed to a single train-test split. It has the ability to remove the variability that might arise as a result of using a single partition of the data for testing. After training 10 previously mentioned models, the “cross_val_score” function from scikit-learn was used to perform a 10-fold cross-validation on the training data [30]. During each iteration, one fold was used as the validation set. The remaining 9 folds were used for training. After each of the 10 iterations, 10 individual accuracy scores were obtained based on how well the models performed on the validation set. The average of these 10 accuracy scores was calculated to get an overall measure of how the models were likely to work on unseen data.

Scikit-learn’s “classification_report” function was used to generate a classification report for evaluating the performance of the ML models [30]. For a given classification problem, it provides an overview of various evaluation metrics for each class. The final report generated by the function includes metrics such as precision, recall, F1-score, and support. The precision metric is a measure of how many of the predicted positive cases were actually correct. A high value of precision corresponds to a low rate of false positives. Recall which is also known as sensitivity or true positive rate is the ratio of correctly predicted positive cases to the total actual positive instances. It is an indication of the model’s ability to recognize all positive cases. Recall value is crucial for ensuring no missing positive instances, even if it means having some degree of false positives. High recall signifies a low level of false negatives. A harmonic mean of precision and recall can be found using the F1-score. Support provides a context about the prevalence of each class and helps understand whether the evaluation metrics are based on a small or large number of instances.

The “confusion_matrix” function provided by the scikit-learn library was used to compute a confusion matrix based on the predictions of the classification models [30]. Confusion matrix is a fundamental tool in ML which is used extensively for measuring the model performance. The main components of a confusion matrix include true positive, true negative, false positive, and false negative.

### Area Under the Receiver Operating Characteristic Curve

In order to quantify the ability of the trained models to distinguish between the positive and negative classes, the concept of the Receiver Operating Characteristic (ROC) curve and its corresponding metric, the Area Under the ROC Curve (AUC ROC) was utilized. For each ML algorithm considered in this study, the AUC ROC score was calculated using the “roc_auc_score” function from scikit-learn [30]. The ROC curve graphically depicts the trade-off between sensitivity and specificity, illustrating how model performance can be tuned by adjusting the classification threshold. The AUC ROC score serves as a comprehensive evaluation metric, capturing the model’s performance across a range of classification thresholds, and providing a single-number summary of its discriminative ability.

### Hyperparameter Tuning

Hyperparameter tuning is a crucial step in optimizing the performance of ML models and was an integral component of the current study. The primary objective of hyperparameter tuning is to identify the optimal configuration of hyperparameters that maximizes the models’ performance. In this study, we utilized the scikit-learn library in Python to conduct comprehensive hyperparameter tuning for multiple ML algorithms. Our hyperparameter tuning process involved utilizing GridSearchCV and RandomizedSearchCV to systematically traverse the hyperparameter space. Each model was evaluated using a 10-fold cross-validation approach, and the selected performance metric, either accuracy or mean squared error, guided the search for optimal hyperparameters. The best-performing hyperparameters were chosen based on the results of the search, ultimately enhancing the generalization ability of our models and ensuring their robustness to overfitting. During the tuning process, we meticulously explored a wide range of hyperparameters summarized in **Table 2**.

**Table 2:** Range of values used for hyperparameter tuning.

| Machine Learning Model | Hyperparamters | Tested Values |
| --- | --- | --- |
| Logistic Regression | C | 0.01, 0.1, 1, 10, 100 |
|  | penalty | 'l2' , 'none'. |
| Support Vector Machine | C | 0.1, 1, 10, 100 |
|  | gamma | 0.1, 0.01, 0.001, 0.0001 |
|  | kernel | 'linear', 'rbf', 'poly' |
| Decision Tree | Max depth | 2, 5, 10, None |
|  | Min samples leaf | 2, 5, 10 |
|  | Min samples split | 1, 2, 4 |
| Random Forest | N estimators | 100, 200, 300, 400, 500 |
|  | Min samples split | 2, 5, 10 |
|  | Min samples leaf | 1, 2, 4 |
|  | Max features | 'sqrt', 'log2' |
|  | Max depth | 5, 10, 15, 20, 25, None |
| Naive Bayes | Var smoothing | 1e-9, 1e-8, 1e-7, 1e-6, 1e-5 |
| K-Nearest Neighbor | N neighbors | 3, 5, 7, 9 |
|  | p | 1, 2 |
|  | weights | 'uniform', 'distance' |
| Gradient Boosting | Learning rate | 0.01, 0.1, 1.0 |
|  | Max depth | 3, 5, 7 |
|  | N estimators | 50, 100, 200 |
| Adaptive Boosting | Learning rate | 0.01, 0.1, 1.0 |
|  | N estimators | 50, 100, 200 |
| Extreme Gradient Boosting | Colsample bytree | 0.7, 0.8, 0.9 |
|  | gamma | 0, 0.1, 0.2 |
|  | Learning rate | 0.1, 0.01, 0.001 |
|  | Max depth | 3, 4, 5 |
|  | N estimators | 100, 500, 1000 |
|  | subsample | 0.7, 0.8, 0.9 |
| Light Gradient Boosting Machine | Learning rate | 0.05, 0.1, 0.2 |
|  | Max depth | -1, 5, 10 |
|  | N estimators | 50, 100, 200 |
|  | Num leaves | 31, 63, 127 |

## Results

### 47,929 Differentially Expressed Genes were Identified

The quality assessment of the raw sequencing data was conducted for a total of 433 raw sequences, revealing that all of them were of high quality. After aligning the reads to the human reference genome, a total of 62,702 genes were discovered. These genes were then subjected to differential expression analysis in the quantification step. A comprehensive analysis revealed that a total of 47,929 genes had differential expression in samples obtained from patients with AD. Among these genes, 10,133 were found to be upregulated, while 5,285 were downregulated. The DEGs were visualized through an MA plot (**Figure 2)**.

**Figure 2:**
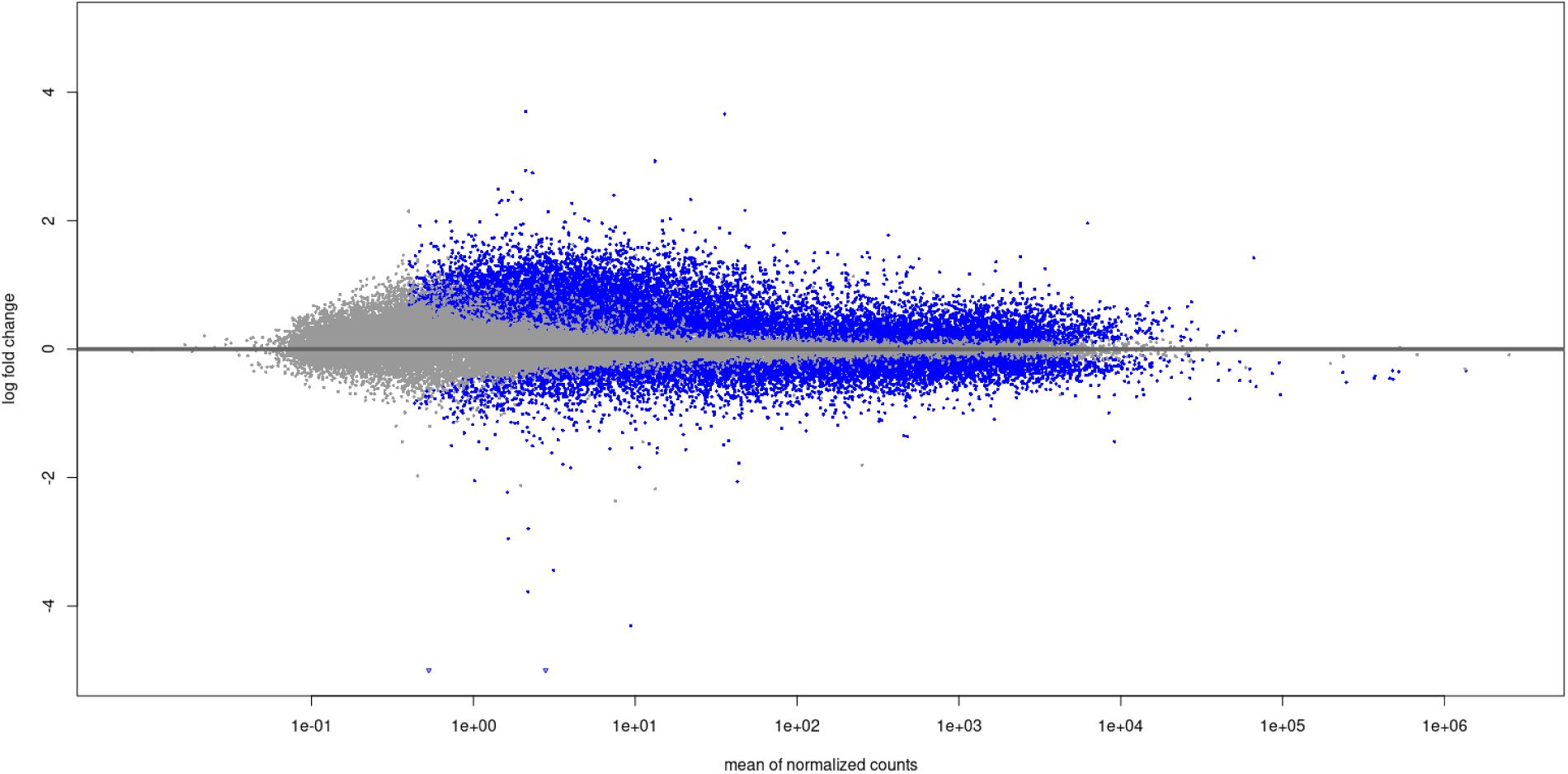
MA plot depicting differentially upregulated and downregulated genes. The upregulated DEGs are displayed at the top of the plot, while the downregulated DEGs are presented at the bottom. The significant DEGs are denoted by blue dots.

### Random Forest Model with Optimized Hyperparameters Exhibited the Best Performance

Top 10 most important features were identified by each of the 4 feature selection tools (**Table 3**). Among these features, 5 genes were found to be commonly identified by all 4 tools (**Figure 3**). These 5 genes (*ITGA10*, *CXCR4*, *ADCYAP1*, *SLC6A12*, and *VGF*) were finally selected as features to be used for training 10 ML models. The input dataset was composed of 433 samples where the number of AD and control samples were 293 and 140 respectively. The train_test_split function split the samples in 80:20 ratio. As a result, 346 samples were used for training the ML models while 87 samples constituted the test dataset. After training and testing the models, their performance was evaluated through metrics such as accuracy, precision, recall, F1-score, AUC ROC, and confusion matrix. After evaluating the models, systematic exploration of a wide range of hyperparameters was carried out in order to find the optimal combination for each of the models. The selected hyperparameter values for the tested models can be found in **Table 4**.

**Figure 3:**
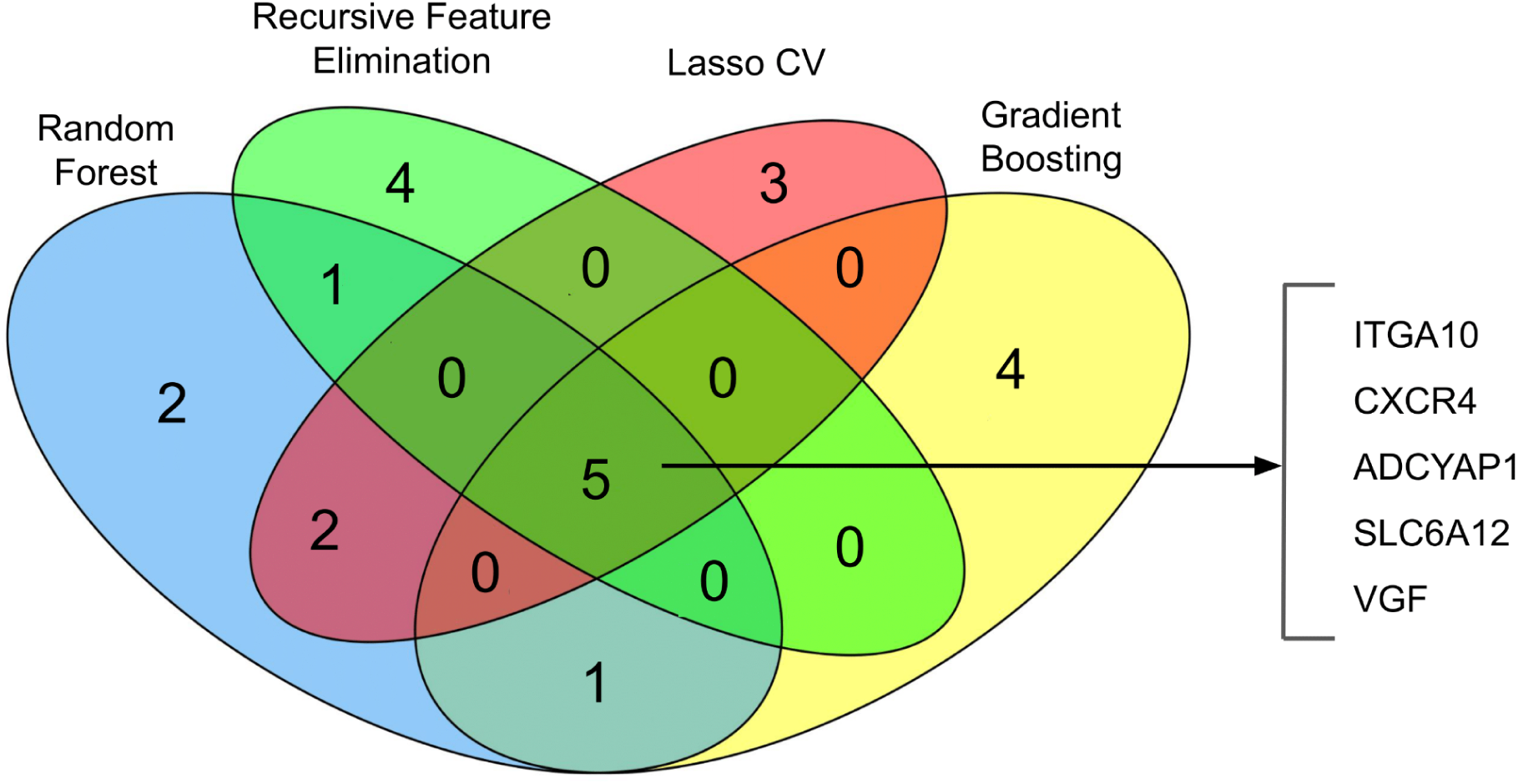
A venn diagram of features (genes) selected by four distinct feature selection algorithms-Random Forest Classifier, Gradient Boosting Classifier, Recursive Feature Elimination, and LassoCV. Five genes were unanimously predicted by all four methods.

**Table 3:** List of feature selection tools and the top 10 most important features.

| Feature Selection Tool | Selected Feature |
| --- | --- |
| Random Forest Classifier | ITGA10, SLC6A12, CXCR4, RIPOR3, KCNE4, ADCYAP1, VGF, ALOX15B, MAS1, S100A4, PCAT19 |
| Gradient Boosting Classifier | ITGA10, CXCR4, SLC6A12, NNAT, VGF, C7orf61, ADCYAP1, RIPOR3, SCARNA7, MTND4P35 |
| Recursive Feature Elimination (RFE) | ITGA10, CXCR4, ADCYAP1, PCAT19, ANKRD36B, RNU1-2, PCDH8, VGF, SLC6A12, MLIP |
| LassoCV | ITGA10, CXCR4, ADCYAP1, SLC6A12, S100A4, VGF, PCDH11Y, DANT2, ALOX15B, H1-4 |

**Table 4:**
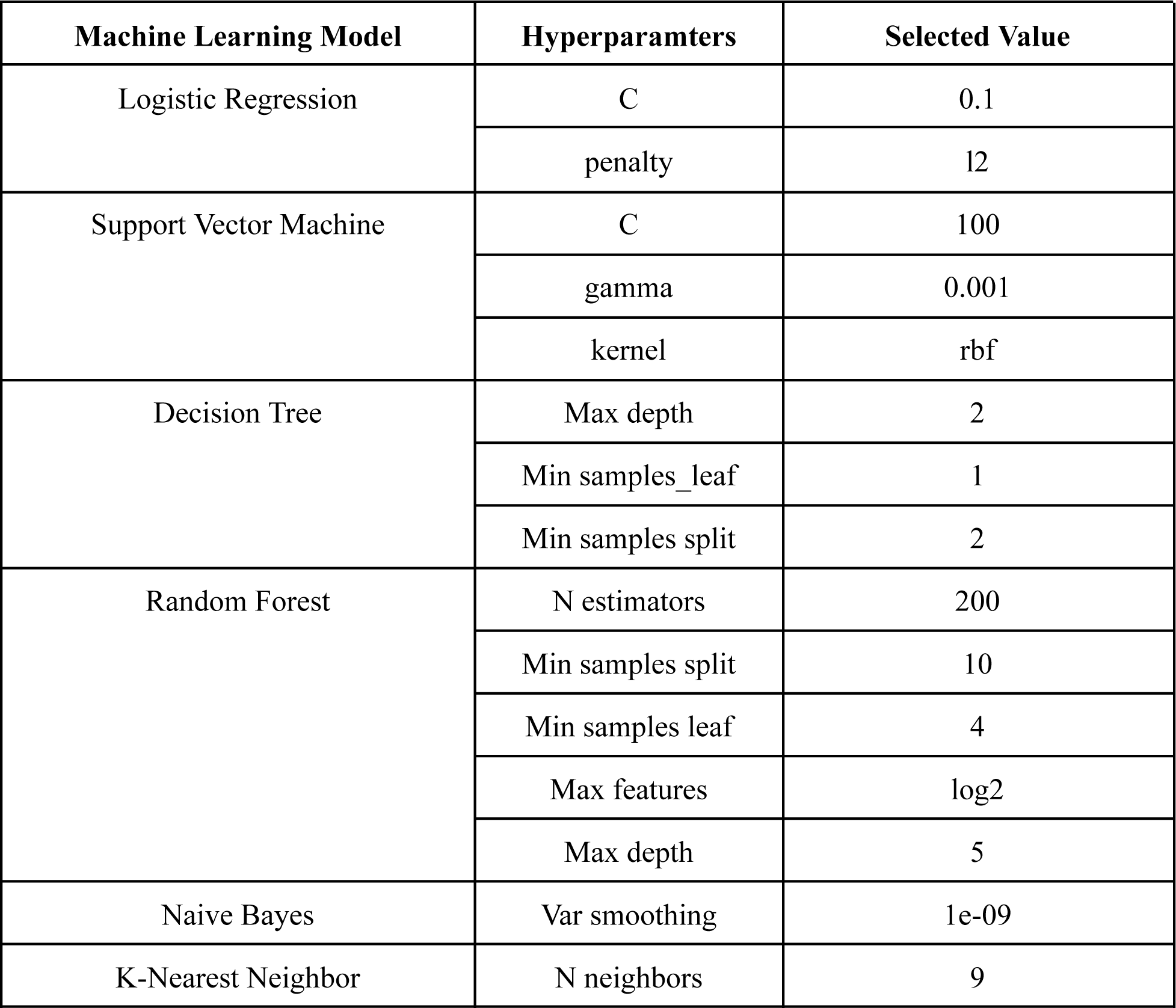
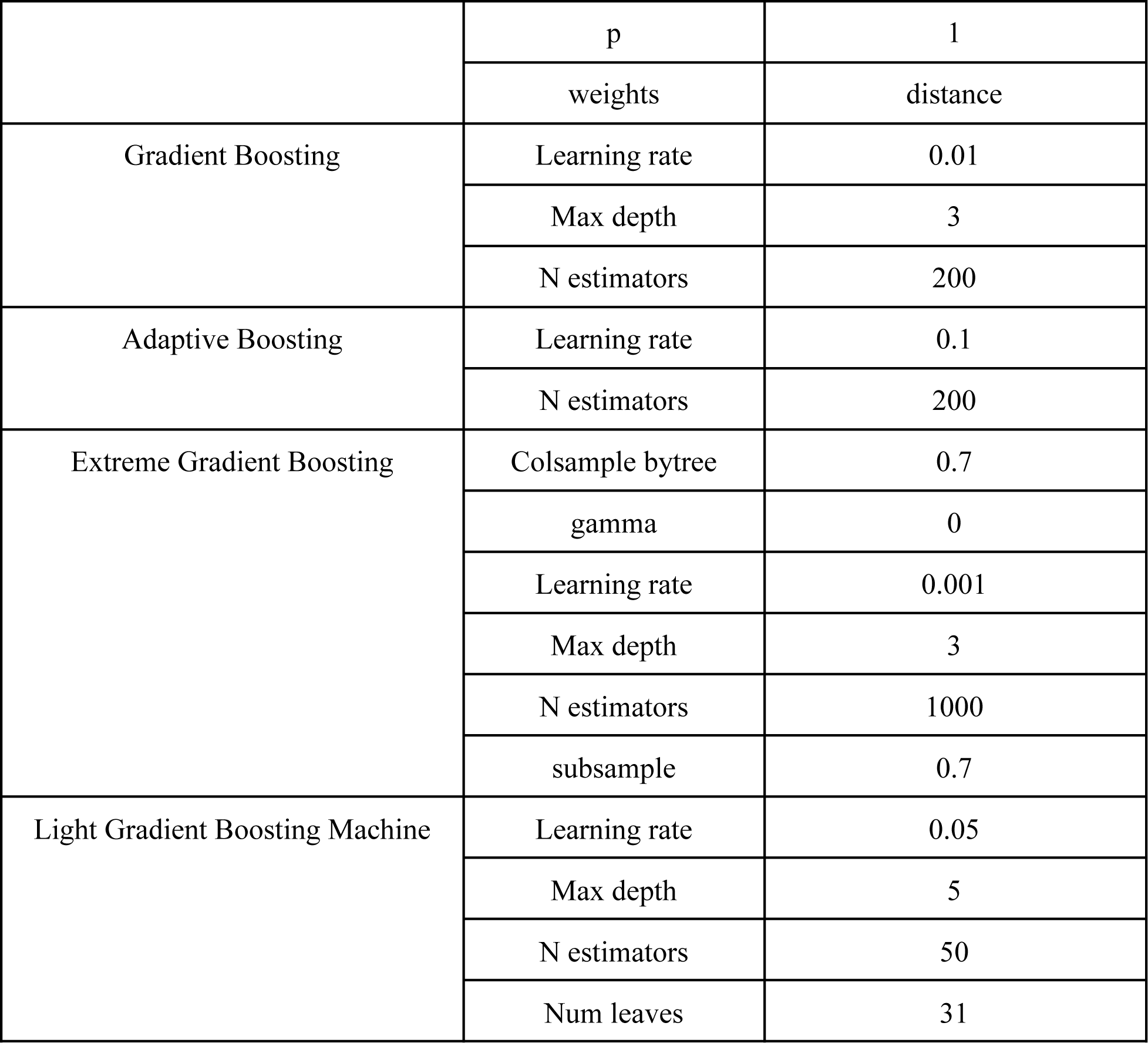
Selected hyperparameter values for machine learning models following tuning.

The accuracy of the models before and after hyperparameter tuning were revealed in **Figure 4**. It depicted that except Logistic Regression and Support Vector Machine for which the accuracy remained the same (0.78), all the models improved their performance following hyperparameter tuning. Before the tuning, the Logistic Regression and Support Vector Machine models had the highest accuracy (0.78) among all. After the tuning, they were surpassed by Random Forest and Gradient Boosting models (0.79). Lowest accuracy was observed for the Decision Tree and K-Nearest Neighbor models both before (0.73) and after (0.75) the hyperparameter tuning.

**Figure 4:**
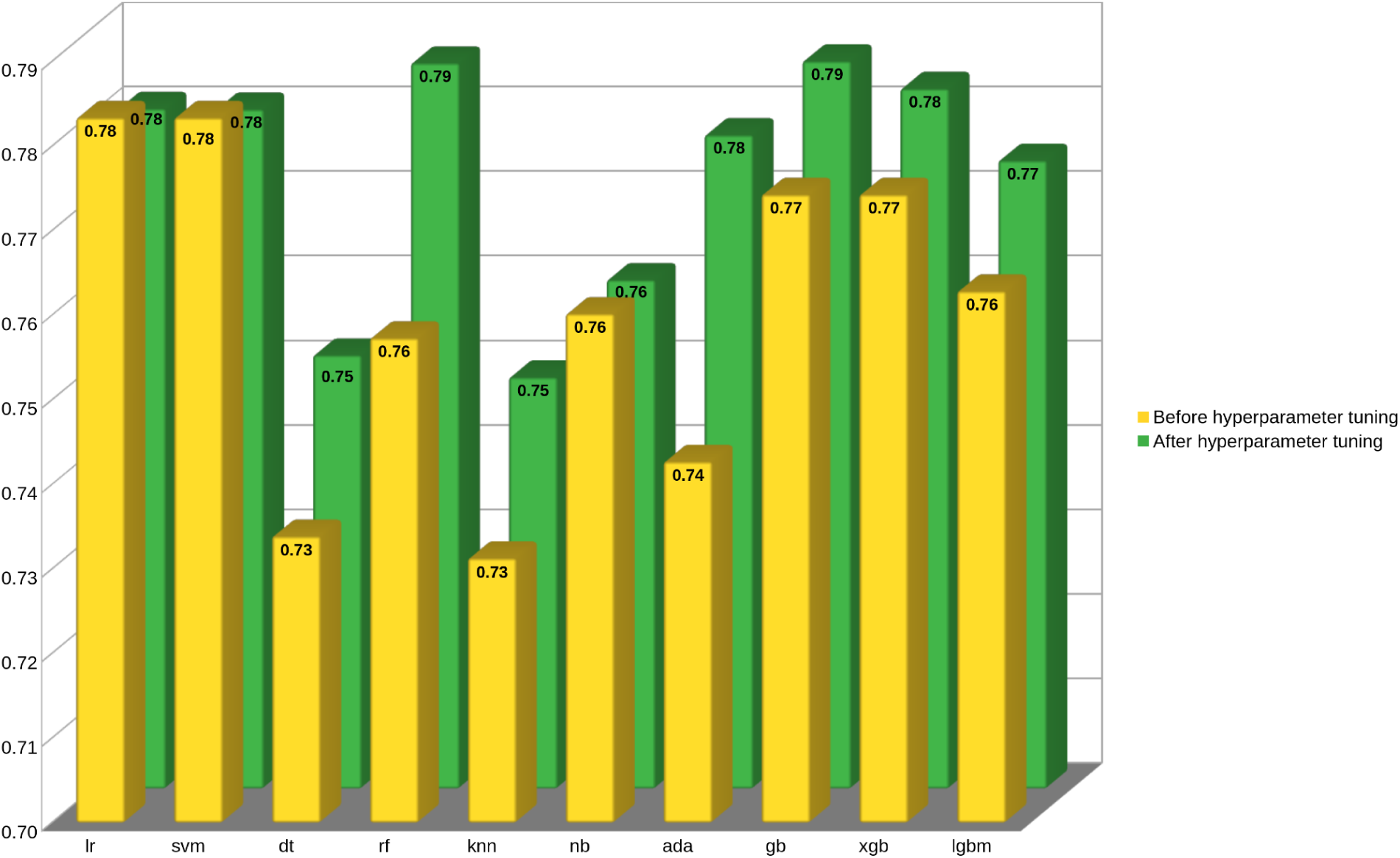
Comparison of accuracy of ML models prior to and following hyperparameter tuning. The accuracy values were calculated after 10-fold cross validation. Yellow bars represent the accuracies before hyperparameter tuning while the green bars represent the accuracies after hyperparameter tuning.

The classification report provided by Scikit-learn’s classification_report function further evaluated the performance of the ML models before and after hyperparameter tuning (**Figure 5**). Random Forest model had the best precision in detecting AD and while the Support Vector Machine had the highest precision for control samples. In terms of recall, the Support Vector Machine possessed the highest value for AD whereas the Random Forest model was found to be the best for control. Support Vector Machine exhibited the highest F1-score for classifying AD samples. However, with respect to controls, the Random Forest model was found to be the best. From the classification report after hyperparameter tuning, highest precision for identifying AD and control samples was observed for Naive Bayes and Logistic Regression models respectively. On the other hand, Logistic Regression and Naive Bayes models had the highest recall values for AD and control respectively. F1-score, which was a balance between precision and recall, revealed that Logistic Regression had the highest value for both AD and control. AUC ROC analysis showed that Naive Bayes had the highest AUC value (0.83) both before and after hyperparameter tuning (**Figure 6**).

**Figure 5:**
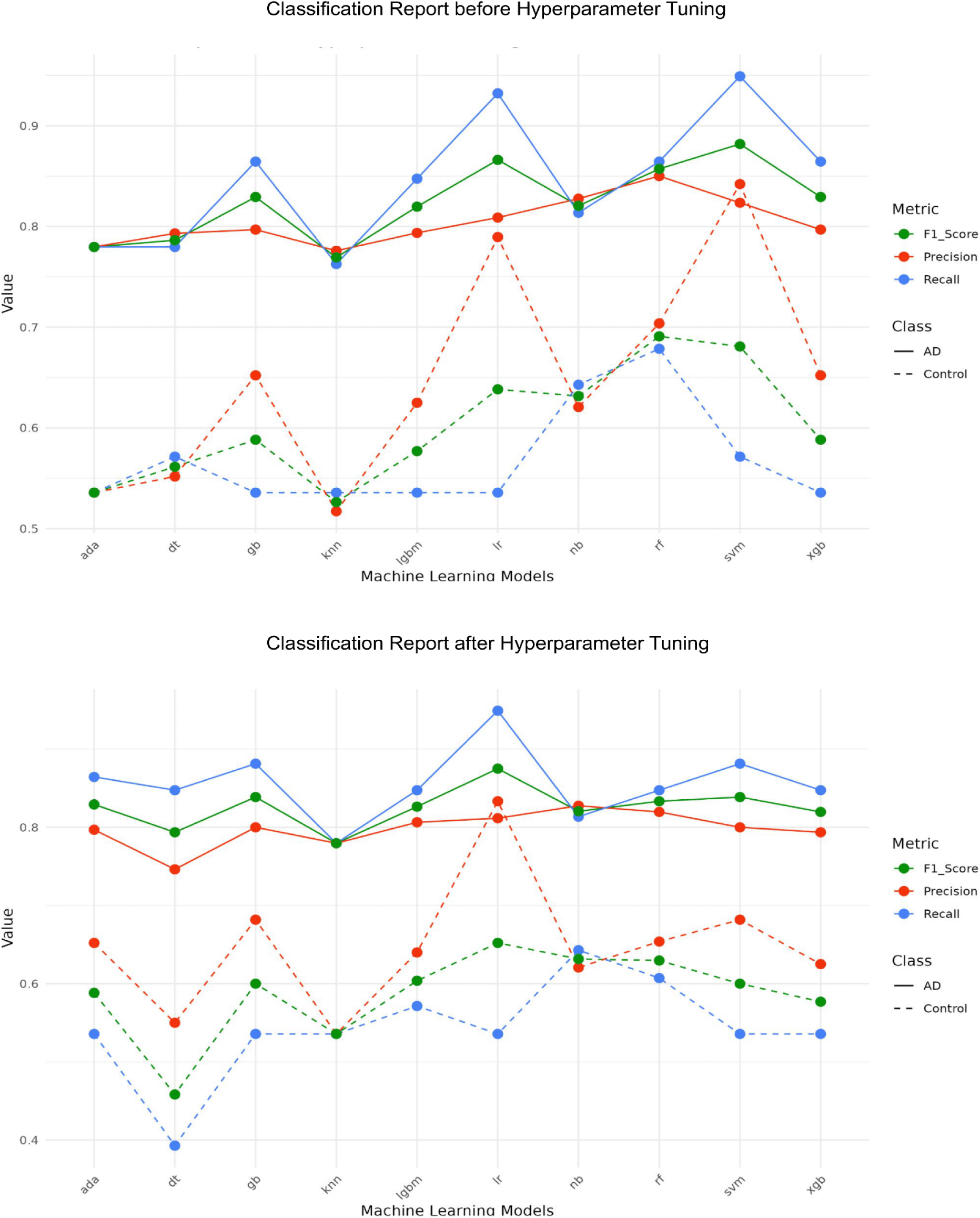
Classification report of the ML models before (top) and after (bottom) hyperparameter tuning. The performance of the models were evaluated based on precision, recall and F1-score. Each metric was measured for both AD and control. The X-axis represents the ML models. The Y-axis represents the values for each metric.

**Figure 6:**
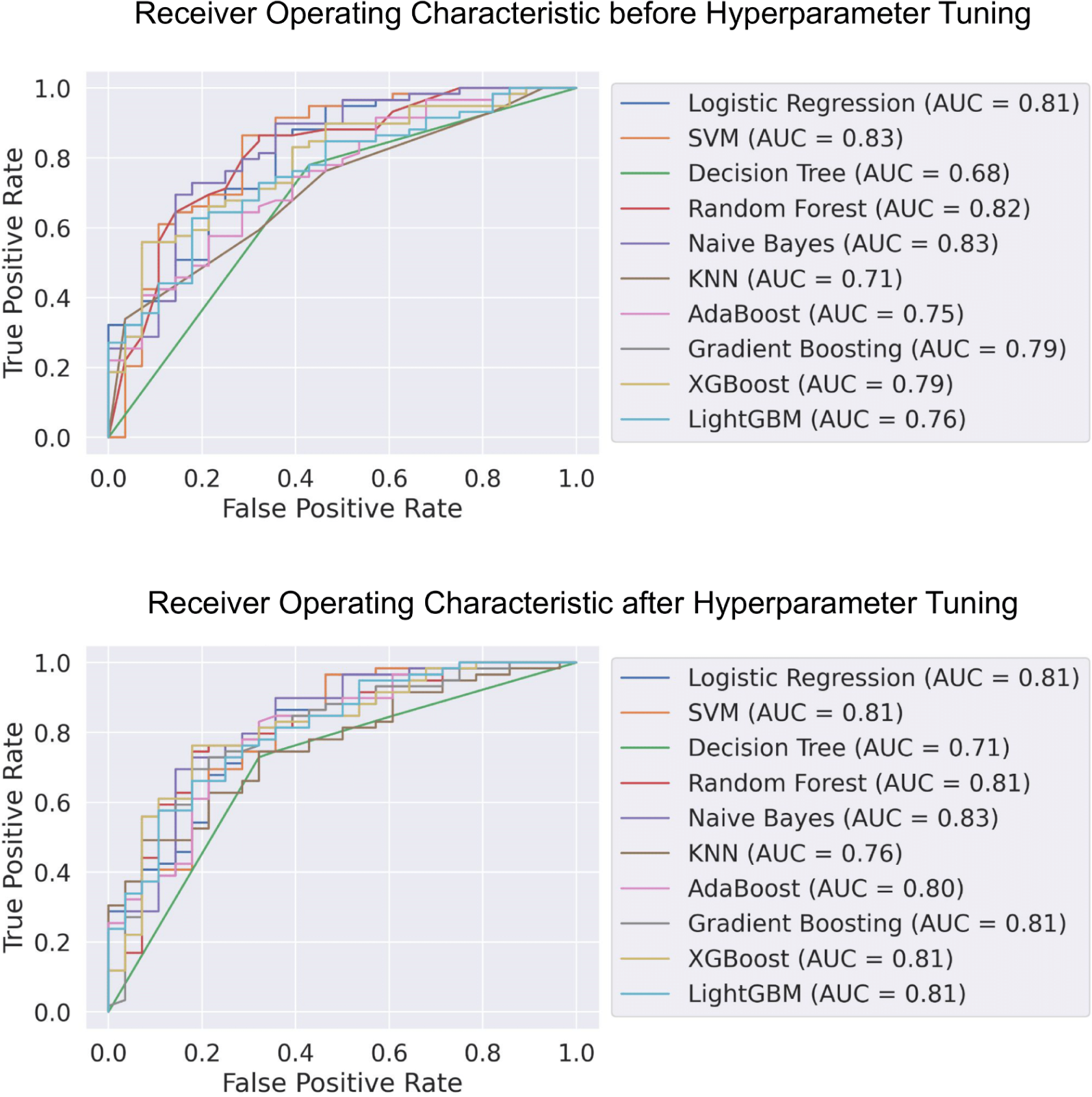
Area under the curve receiver operating characteristic of the models before (top) and after (bottom) hyperparameter tuning. The X-axis represents the false positive rate while the Y-axis represents the true positive rate.

In order to visualize how many of the predictions on the test data (n= 87) were correct before and after hyperparameter tuning, confusion matrix diagrams were generated and the results were depicted in **Figure 7**. The diagram showed that the highest number AD samples were correctly identified as AD (n=54) by the Gradient Boosting model before hyperparameter tuning. However, Adaptive Boosting had the highest number of accurate predictions (n=52) in terms of AD after the tuning. While predicting the control as control before the hyperparameter tuning, Support Vector Machine was the best since it correctly identified 18 control samples as controls. After the tuning, Random Forest and Extreme Gradient Boosting models exhibited the best performance in this regard by identifying 18 control samples correctly.

**Figure 7.**
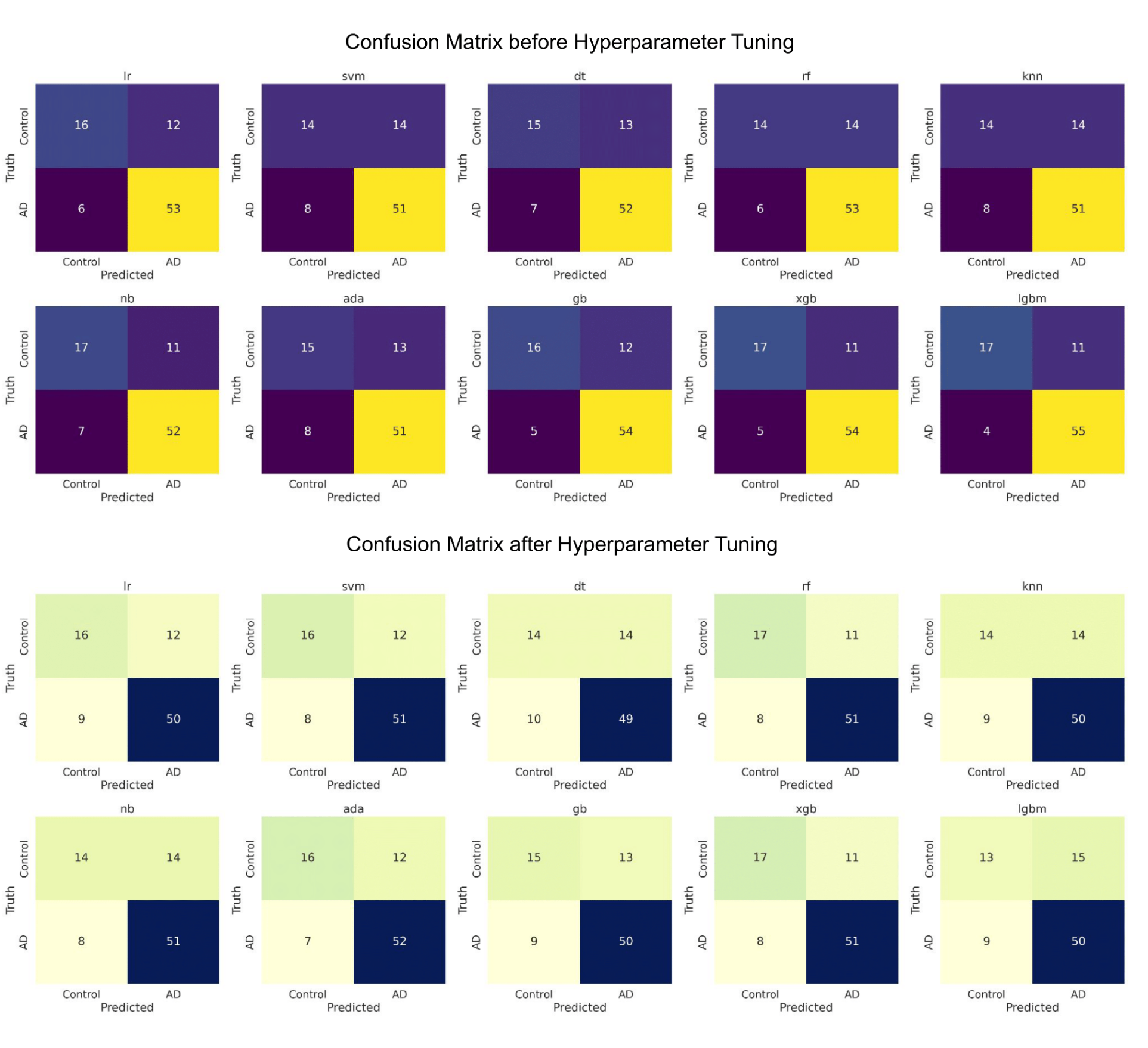
Confusion matrix of the machine learning models before (top) and after (bottom) hyperparameter tuning. The X-axis depicts the predictions while the Y-axis depicts the truth.

Based on all the metrics, the Random Forest model with specific hyperparameters (N estimators=200, Min samples split=10, Min samples leaf= 4, Max features= log2, Max depth= 5) was determined as the best model among all for our purpose. It had the highest accuracy (0.79), second highest AUC value (0.81), third highest F1 score, second highest number of accurate predictions of AD as AD and highest number of predictions of control as control. This cutomized random forest model was utilized to develop the Alzheimer’s Identification Tool using RNA-Seq (AITeQ) which can be implemented through google collaborator. The full AITeQ documentation containing the instructions on how to run the tool for AD prediction can be found at https://github.com/ishtiaque-ahammad/AITeQ.

## Discussion

Integration of ML methods with transcriptomics data processing has been reported to benefit the understanding of complicated neurodegenerative illnesses like AD. Along these lines, the current study aimed at analyzing RNA-Seq data using ML algorithms to predict AD. The findings from this study will contribute to the ongoing efforts for early and precise diagnosis of AD by utilizing a refined 5-gene signature as an accurate predictor of the disease.

The work relied heavily on the thorough analysis of RNA-Seq data from publicly available datasets in NCBI. Quality evaluation, read alignment, and quantification constituted parts of the preprocessing processes essential for generating valid inputs for the ML models in the subsequent step. The complex transcriptomic aberrations associated with AD were highlighted by the finding that over 47,000 genes undergo differential expression in individuals with the condition. However, after applying a stringent filtering scheme to identify the genes with the most significantly altered expression, only 59 of them remained.

One of the most crucial aspects of this study was the selection of features (genes) while developing a robust predictive model for AD. Using a combination of techniques, such as the Random Forest Classifier, Gradient Boosting Classifier, Recursive Feature Elimination, and LassoCV, 5 genes were consistently determined to be important across all employed techniques. Implementing multiple methods improved the credibility of the gene signature, resulting in a more dependable method for predicting AD. This was in accordance with a number of previously conducted ML studies that utilized feature selection to solve classification problems more accurately especially in the biological domain [31], [32], [33], [34].

ML algorithms formed the basis of the predictions made in this investigation. The flexibility and power of ML was characterized by the use of a wide variety of algorithms to recognize patterns from RNA-Seq data. These included Logistic Regression, Support Vector Machines, and Ensemble approaches like Random Forest and Gradient Boosting. Following the example of other published studies that systematically experimented with hyperparameters led to improved model performance in our study [35], [36].

A sign of the intricacy of AD categorization is the discovery of trade-offs across various measures (Accuracy, precision, recall, F1-score, and AUC ROC) used to evaluate the model performance. This is a common practice followed by a number of earlier studies that have emphasized the necessity to use multiple criteria to objectively evaluate classification models instead of relying on a single one [37], [38], [39].

It is to be noted that the most promising result of this study is the establishment of a 5-gene signature that holds true across all ML models. This signature has the potential to be integrated into a biomarker panel for AD diagnosis. There is a history of such gene signature based novel diagnostic biomarkers discovery using integrated ML and transcriptomic investigations for example-in cases of AD [40], breast cancer [37], COVID-19 [41], psoriasis [42], tuberculosis [43] and so on.

The set of 5 genes (*ITGA10, CXCR4, ADCYAP1, SLC6A12, VGF*) identified through our investigation demand closer attention in terms of their relationship with AD. Interestingly, *ITGA10* and *SLC6A12* were also suggested as potential diagnostic biomarkers in a study published in 2023 that compared the expression profiles of AD and control tissue samples using ML algorithms [44]. *CXCR4* is a highly conserved seven-transmembrane G-protein-coupled receptor that has been reported to be involved in the pathophysiology of AD [45]. Neuronal expression of *ADCYAP1* has been observed to be significantly reduced throughout the brain of AD patients [46]. *VGF* has been suggested as a biomarker and a therapeutic target in neurodegenerative diseases [47]. Hence, all 5 genes detected in our study as a crucial set of predictors of AD have been reported to be associated with AD in one way or the other.

While translating the findings from this study, there are a number of caveats to keep in mind despite the encouraging results. It is a retrospective study that relies on a limited number of datasets. As a result, it has the potential to introduce certain biases that might impact how well the conclusions generalize to new data. Therefore, it is imperative to validate the findings with a larger number of samples collected from a variety of demographics. Another challenge is the variability in RNA-Seq data due to biological and technical factors such as batch effects, sequencing depth, and normalization methods. Batch effects can lead to spurious correlations between genes and disease outcomes, while sequencing depth and normalization methods can affect the accuracy and reproducibility of gene expression measurements [48]. ML algorithms can be sensitive to these factors, and appropriate data preprocessing and normalization methods are necessary to ensure accurate classification results [49]. Apart from all these, the multifaceted nature of neurodegenerative disorders, including but not limited to non-coding RNA-mediated regulations, protein-protein interaction networks, and epigenetic alterations, calls for an approach that goes beyond just focusing on gene expression.

## Conclusion

Results from the current study opened up several promising new lines of inquiry. The promise of ML in understanding the complex nature of AD has been demonstrated by its application on disease prediction from RNA-Seq data. The importance of a possible biomarker panel for accurate diagnosis of AD is highlighted by the discovery of a consistent 5-gene signature. It is crucial to further investigate the functional role played by the identified 5-gene signature with respect to AD etiology. The diagnostic potential of the gene signature should be validated in subsequent studies involving a variety of populations through longitudinal investigations.

## Author Contributions

I.A., A.B.L, A.B. and M.S.A. trained and evaluated the machine learning models and developed the AITeQ framework. A.B.L and T.B.J. conducted the differential gene expression analysis and data preprocessing for the machine learning models. I.A., A.B.L. wrote the original draft of the manuscript. Z.M.C., M.U.H., and K.C.D. reviewed and edited the manuscript. M.S. and C.A.K. supervised the research project.

## Data Availability

The data generated in this study are included within the manuscript and the supplementary files.

## Code Availability

Code for using the AITeQ model is available at: https://github.com/ishtiaque-ahammad/AITeQ.

## Funding

This research work did not receive any funding.

## Competing interests

The authors declare no competing interests.

### Author Biographies

### Ishtiaque Ahammad

Ishtiaque Ahammad is a Scientific Officer at the Bioinformatics Division, National Institute of Biotechnology.

### Anika Bushra Lamisa

Anika Bushra Lamisa is a Post-Graduate Research Fellow at the Bioinformatics Division, National Institute of Biotechnology.

### Arittra Bhattacharjee

Arittra Bhattacharjee is a Scientific Officer at the Bioinformatics Division, National Institute of Biotechnology.

### Tabassum Binte Jamal

Tabassum Binte Jamal is a Post-Graduate Research Fellow at the Bioinformatics Division, National Institute of Biotechnology.

### Md. Shamsul Arefin

Md. Shamsul Arefin is a graduate student at the Department of Biochemistry and Microbiology, North South University.

### Zeshan Mahmud Chowdhury

Zeshan Mahmud Chowdhury is a Scientific Officer at the Bioinformatics Division, National Institute of Biotechnology.

### Mohammad Uzzal Hossain

Mohammad Uzzal Hossain is a Scientific Officer at the Bioinformatics Division, National Institute of Biotechnology.

### Keshob Chandra Das

Keshob Chandra Das is a Chief Scientific Officer (C.C.) at the Molecular Biotechnology Division, National Institute of Biotechnology.

### Dr. Chaman Ara Keya

Dr. Chaman Ara Keya is an Associate Professor at the Department of Biochemistry and Microbiology, North South University.

### Dr. Md Salimullah

Dr. Md Salimullah is the Director General (A.C.) and Chief Scientific Officer at the National Institute of Biotechnology.

## Notes

https://github.com/ishtiaque-ahammad/AITeQ

### Competing Interest Statement

The authors have declared no competing interest.

